# The effect of dietary intervention on inflammatory markers in patients with type 2 diabetes. An analysis of the Lifestyle Over and Above Drugs in Diabetes (LOADD) study

**DOI:** 10.1101/483412

**Authors:** Chris S Booker, Kirsten J Coppell, Ashley Duncan, Minako Kataoka, Sheila M Williams, Jim I Mann

## Abstract

Ninety-three participants with poorly controlled type 2 diabetes despite optimized drug therapy were randomised to receive intensive dietary advice or usual care for six months. Following dietary intervention a significant reduction in interleukin-18 levels was observed, with a ratio of change (95% CI) of 0.90 (0.82-0.99); p = 0.033. Changes in IL-18 correlated with changes in neopterin, r = 0.299 (p = 0.009)

## 1. Introduction

There is increasing recognition that obesity and type 2 diabetes are characterized by sub-clinical inflammation, which may contribute to the development of atherosclerosis [1]. Since intensive glycaemic control has not been found to reduce cardiovascular events in people with type 2 diabetes [2], reducing inflammation may be an important focus for reducing cardiovascular morbidity and mortality in this group of patients. Diabetes lifestyle intervention aims to improve glycaemic control, lipids and blood pressure levels, but appropriate dietary changes may also influence levels of inflammation. We therefore measured a selection of inflammatory markers in the Lifestyle Over and Above Drugs in Diabetes (LOADD) study, a randomized controlled trial which investigated the effect of a six month intensive dietary intervention in 93 participants with type 2 diabetes with poor glycaemic control whose drug therapy had been optimized according to treatment guidelines [3].

## 2. Material and methods

### 2.1 Study design and previous findings

The LOADD study has been described in detail elsewhere [4]. The study was approved by the Lower South Regional Ethics Committee of New Zealand (LRS/05/07/026), and registered prior to commencement (NCT00124553). Participants were randomized to an intensive evidence-based dietary intervention group, which received advice based on recommendations from the European Association for the Study of Diabetes [5], or a usual care control group. Clinically and statistically significant improvements (95% CI) in weight of 1.3 kg (0.1, 2.4), waist circumference of 1.6 cm (0.5, 2.7), and HbA_1c_ of 0.4% (0.1, 0.7) [4 mmol/mol (1, 8)] were observed in the intervention group [4].

### 2.2 Sample processing and laboratory measurements

CRP was measured using a high sensitivity CRP kit (Roche, catalogue number 04628918190) on a Cobas c311 analyser; participants with CRP values > 10 mg/L were excluded from further analysis. IL-18 was measured by ELISA (MBL, Japan); leptin and neopterin by ^125^I-radioimmunoassay (Leptin: Millipore, MA, USA, catalogue number HL81K; Neopterin: BRAHMS Diagnostica, Germany, BRAHMS Neopterin RIA); IL-18BP using a commercial IL-18BPa ELISA (RayBiotech, Inc., GA, USA, catalogue number ELH-IL-18 BPA-001). Remaining cytokines were measured by Lumenix™ bead array multiplex system using beads and plates from Millipore (MA, USA) and a Bio-Plex suspension array instrument (Bio-Rad Laboratories, CA, USA). Participant samples were analysed in duplicate in RIAs and ELISAs, and using single samples in multiplex and CRP assays. Samples from RIAs and ELISAs with coefficients of variation ≥ 20% were excluded from further analysis.

### 2.3 Statistical analysis

The data were analysed according to modified intention to treat. Multiple regression was used to estimate the difference between treatments adjusting for the baseline, age and sex. As the data were log transformed before analysis the differences are presented as the ratio (95% confidence interval) of the geometric means at six months. Pairwise correlation coefficients to assess the relationship between changes in IL-18 and neopterin were generated using non-log transformed measures from all participants.

## 3. Results

The effect of the LOADD study dietary intervention on levels of adipokines and inflammatory markers is shown in Table 1. The intervention group exhibited significant reductions in IL-18 and neopterin levels relative to the control group. No significant changes were observed in other inflammatory markers and adipokines, although neopterin levels showed a decrease of borderline significance (p = 0.057). After adjustment for baseline BMI and HbA_1c_, the effect size for the change in IL-18 was minimally altered, with a difference of 0.90 (95% CI: 0.82 – 0.99, p=0.033). Changes in IL-18 and neopterin were correlated, with a pairwise correlation coefficient of r = 0.299 (p = 0.009, n = 76).

**Table 1:** The effect of usual dietary advice and intensive evidence based dietary advice intervention on adipokines and inflammatory markers in patients with type 2 diabetes and poor glycaemic control despite optimized drug therapy.

|  |  | Intervention |  | Control |  | Difference |  |
| --- | --- | --- | --- | --- | --- | --- | --- |
| Measure | n | baseline * | 6 months * | baseline * | 6 months * | (95 % CI) | p |
| CRP (mg/L) | 41 | 1.9 (1.1, 3.7) | 1.4 (0.7, 3.2) | 2.3 (0.9, 3.5) | 2.4 (0.9, 3.9) | 0.76 (0.53, 1.07) | 0.118 |
| IL-6 (pg/ml) | 29 | 4.94 (1.43, 9.76) | 5.61 (1.88, 23.21) | 4.68 (1.64, 10.32) | 4.44 (1.78, 20.30) | 1.05 (0.47, 2.32) | 0.906 |
| IL-10 (pg/ml) | 20 | 3.08 (1.74, 6.61) | 2.42 (1.60, 8.63) | 3.81 (2.01, 5.27) | 5.42 (2.39, 11.96) | 0.58 (0.29, 1.16) | 0.120 |
| IL-18 (pg/ml) | 45 | 506 (430, 666) | 449 (354, 553) | 527 (418, 657) | 504 (419, 652) | 0.91 (0.83, 1.00) | 0.049 |
| IL-18 BP (pg/ml) | 32 | 455 (140, 857) | 362 (163, 768) | 473 (255, 751) | 426 (229, 782) | 0.94 (0.62, 1.44) | 0.783 |
| Leptin (ng/ml) | 36 | 36.4 (28.8, 48.0) | 37.4 (20.9, 54.5) | 30.2 (14.7, 45.6) | 33.2 (12.8, 51.6) | 1.12 (0.83, 1.52) | 0.445 |
| Neopterin (nmol/l) | 39 | 5.36 (4.55, 6.09) | 5.04 (4.42, 5.47) | 5.08 (4.51, 5.83) | 5.21 (4.56, 5.85) | 0.89 (0.79, 1.00) | 0.057 |
| sTNF-R1 (pg/ml) | 40 | 836 (615, 1019) | 901 (671, 1127) | 857 (611, 960) | 854 (661, 1093) | 1.00 (0.89, 1.12) | 0.964 |
| sTNF-R2 (pg/ml) | 40 | 2727 (2413, 3300) | 2919 (2418, 3254) | 2616 (2155, 3157) | 2648 (1908, 3613) | 1.09 (0.85, 1.39) | 0.482 |
| TNFα (pg/ml) | 44 | 9.7 (6.7, 16.3) | 14.0 (10.2, 21.8) | 10.0 (6.7, 13.9) | 15.0 (12.0, 18.9) | 1.01 (0.79, 1.29) | 0.938 |

\* Values are median (IQR)

## 4. Discussion

This randomized controlled trial examined the effects of an evidence-based six month dietary intervention in patients with type 2 diabetes on a range of adipokines and inflammatory markers. The intervention group exhibited a significant reduction in circulating IL-18 levels relative to the control group, an inflammatory marker which may have an active role in the development of cardiovascular disease [6,7]. These reductions remained significant after further adjustment for baseline BMI and HbA_1c_ levels. While obesity is associated with increased inflammation, these results suggest that an evidence-based dietary intervention can lower select inflammatory markers independent of adiposity and glycaemic control

A reduction in neopterin levels of borderline significance was also observed. Neopterin release from macrophages is induced by IL-18 (via induction of IFN-γ [8]), and increases in circulating neopterin levels have been observed in monkeys after an injection of IL-18 [9]. It is likely that changes in neopterin therefore occurred downstream of the reduction in circulating IL-18 levels in our study, as changes in neopterin levels were correlated with changes in IL-18.

### 4.1 Comparisons with previous research

The LOADD study dietary intervention resulted in a significant increase in the percentage of total energy (%TE) consumed as protein, a significant reduction in %TE consumed as saturated fat, and an increase in dietary fibre intake of borderline significance [4]. Previous metabolic studies by Esposito *et al*. have shown that IL-18 increases after a high fat meal and decreases after a high fibre meal [10]. Changes in inflammatory markers in the LOADD study are therefore consistent with the short-term changes observed by Esposito *et al.* [10]. The mechanism underlying these changes is unclear, but there is evidence that dietary modification influences gut microbiota [11], that IL-18 is produced in the gut [12], and that different bacteria produce diverging effects on IL-18 expression in intestinal cells *in vitro* [13]. Therefore our observations could be explained by the interaction of dietary components with gut bacteria resulting in altered levels of circulating inflammatory cytokines.

### 4.2 Study limitations

Chance may explain the significant reduction in IL-18 levels observed in our study. However, changes in neopterin levels are induced by IL-18, and the correlated changes in neopterin and IL-18 suggest the significant reduction in IL-18 is likely to reflect a true biological effect of the intervention.

### 4.3 Conclusions

The relationship between an evidence-based dietary intervention and reduction in IL-18 levels, independent of adiposity and glycaemia, suggests a further mechanism by which diet may influence cardiovascular risk in people with type 2 diabetes.

## Acknowledgements

We thank the participants of the LOADD study. Many thanks to Michelle Harper and Holiday Wilson, Diabetes and Lipid Laboratory, Department of Human Nutrition, University of Otago for technical assistance.

## Author contributions

KJC co-conceived the main study; contributed to the design, implementation, conduct, and monitoring of the main study; and interpretation of results and editing this manuscript. JIM co-conceived the main study; contributed to the design, implementation, conduct, and monitoring of the main study; analysis and interpretation of results, and was responsible for editing the manuscript. MK participated in the implementation and conduct of the dietary intervention and contributed to the analysis and interpretation of diet records. AD and CSB were responsible for conducting the laboratory analyses. SMW contributed to the design; performed the statistical analyses; and contributed to the interpretation of the findings. CSB contributed to the design of the study and was responsible for writing the manuscript drafts. All authors approve of the final manuscript and decision to publish.

## Abbreviations

CRP: C-reactive protein
IL-6/10/18: interleukins 6 10 or 18
IL-18BP: interleukin-18 binding protein
TNFα: tumour necrosis factor alpha
sTNF-R1/2: soluble tumour necrosis factor receptors 1 or 2
IFN-γ: interferon gamma

